# DRUGSYNC: Prediction of Synergistic Drug Combinations Using GCN Graph Convolutional Network based and Pre-learned Drug-induced Gene Expression Profiles

**DOI:** 10.1101/2024.12.05.627115

**Authors:** Longfei Zhao, Jie Luo

## Abstract

Drug-drug synergy (DDS) is crucial for identifying novel and effective drug combinations. With the accumulation of experimental data (DrugCombDB) and the uprising of deep learning approaches, many efforts have been made to computationally predict effective drug pairs. However, most models had limited success, with the highest ROC AUC of 0.65 for predicting pairs of unseen drugs. Drug-induced gene expressions profiles (DGEP) from datasets like LINCS L1000 offer a robust proxy to describe drug effects. However, overlap between DrugCombDB and experimental DGEP data is sparse only ∼2%. To address these limitations, we propose the DRUGSYNC (**<u>D</u>**rug **<u>R</u>**esponse and **<u>U</u>**tilization for **<u>G</u>**raph-based **<u>SYN</u>**ergy **<u>C</u>**lassification) prediction framework that leverages pre-learned drug-target interaction (DTI) and pre-learned DGEP as features. Graph Convolutional Network (GCN) was used to represent the inherent network of genetic interactions. Our pre-learned DGEP achieved 0.877 Pearson correlation coefficients (PCC). Furthermore, our framework excels by integrating pre-learned DGEP and DTI of paired drugs for DDS prediction, achieving superior performance over benchmark technologies. Notably, it demonstrates exceptional accuracy in predicting interactions between unseen drugs, with an ROC AUC score of 0.854. The analysis further emphasized the importance of accurate DGEP predictions. This method can be extended to any cell lines.

## Introduction

Combination therapy with multiple drugs has been shown to significantly improve therapeutic efficacy, and address limitations of monotherapy, such as reducing drug resistance and toxicity [1,2]. As a result, drug combination therapy has become a cornerstone the primary treatment strategy to manage for many complex diseases [3]. However, the exponential number of possible drug pair combinations make experimental testing prohibitively expensive.

Recent advances in artificial intelligence (AI) technology have enabled novel prediction methods for DDS, aiming to expand the virtual search space of combination therapy while reducing experimental costs [4]. Databases such as DrugComb [5,6], DrugCombDB [7], SYNERGxDB [8], NCI-ALMANAC [9], O’Neil [10] et al. study and AZ-DREAM mainly focus on anti-cancer drug combinations are the most popular testbeds for new algorithms and softwares for DDS. DeepSynergy utilized molecular fingerprints to predict DDS [11]. DeepDDS utilizes RDKit to transform the SMILES notation into molecular graphs, subsequently employing either a GCN or a Graph Attention Network (GAT) to compute the drug embedding vectors [12]. DFFNDDS transforms SMILES notation into vectors utilizing a fine-tuned BERT model, which is a machine learning framework designed for natural language processing [13]. Most of these methods focus on the chemical descriptors of drugs and gene expression profiles of cell lines rather than their interaction. GraphSynergy leverages the proteins modules of drugs in the protein-protein interaction (PPI) network from prior knowledge [14]. KGANSynergy is a novel knowledge graph attention network to predict drug synergy, which utilizes heterogeneous graph information of known drug–protein association, cell line–protein association, protein–protein association, and cell line–tissue association effectively [15]. However, these models suffer from a significant limitation: many studies evaluate model performance using leave-combinations-out cross validation (CV), reaching as high as ROC AUC 0.97 [16], but failed to report their predictive performance on either novel drugs or new cell lines [11–15]. Recent works start to evaluate model with leave-drugs-out CV, and leave-cells-out CV, revealing much lower predictive accuracy. The challenge can be attributed to the following reasons: (i) Information leakage when evaluating models with leave-combinations-out sparadigm; (ii) A limited sample size, for works using some of the early databases, leading to possible overfit; (iii) An inadequate representation of drug-cell line interactions, resulting in the failure to capture important feature that may contribute to predictions.

Previous studies have demonstrated the value of DGEP as characteristics of drugs for predicting the efficacy and importance of drug-drug interaction [17–21]. Cell line-specific DGEP is arguably the best proxy to describe interaction between drug and cell lines. Lin et al. [20], and Han et al. [21] add the cell line-specific DGEP as a new feature type to improve the accuracy of DDS prediction. Even low accuracy prediction of DGEP (PCC = 0.5) has aided in improving DDS model performance. Moreover, how the drug-induced gene expression profile of a single drug specifically influences drug interactions remains a complex issue [22,23]. Utilizing the concept of interaction regulation networks might be an effective approach to this problem [24]. The network structure largely determines the dynamics of the interacting molecules, hence the function it can fulfill. Drug interactions may also be determined in such a manner, so that the structure of the biological network involving the drug targets under study may shed light into the way the drugs act and interact [25–27]. These relationships primarily encompass the modulation of target proteins by drugs and the interactions between proteins.

It is plausible that building on curation of drug synergy pairs in DrugCombDB, and the high quality DGEP from datasets like LINCS L1000 [28], as well as proper feature representation, one can significantly improve DDS prediction. However, only 1% drug pairs in DrugCombDB have available experimental DGEP. To this end, we propose to (i) expand number of DGEP for drugs with at least one experimental DGEP to any cell line, using deep neural network (DNN) learning with high accuracy; (ii) to construct a model based graph convolutional representation using protein-protein interaction (PPI) networks; (iii) to incorporate drug-target interaction (DTI) information as key features in a machine learning framework designed to predict synergistic drug combinatorial efficacy. We then test the model performance with both classification and regression tasks, using leave-combinations-out, leave-drugs-out, as well as leave-cells-out paradigms.

## Results

### 1. Overview

Our proposed model framework is composed of three modules designed to predict DDS. Firstly, we constructed a predictive model based on the LINCS L1000 project to obtain high quality DGEP. By enhancing the volume of high-quality data in the LINCS L1000, this process significantly increases the overlap between LINCS L1000 and DrugCombDB datasets. Secondly, we constructed a DTI matrix integrating our proprietary MVAE-DFDPnet predictive models [27] with the extensive data available in the BindingDB database [29]. The DTI matrix was designed to capture comprehensive interaction profiles between drugs and genes. Finally, we employed a PPI network and GCN to extract graph representation of DGEP. In this model, the features of DGEP and DTI of two paired drugs were mapped onto the nodes of the PPI graph. Through multiple layers of graph convolutions, the model learned the interactions between drugs and gene expressions. Ultimately, the model outputs a synergy score (Fig. 1).

**Fig. 1.**
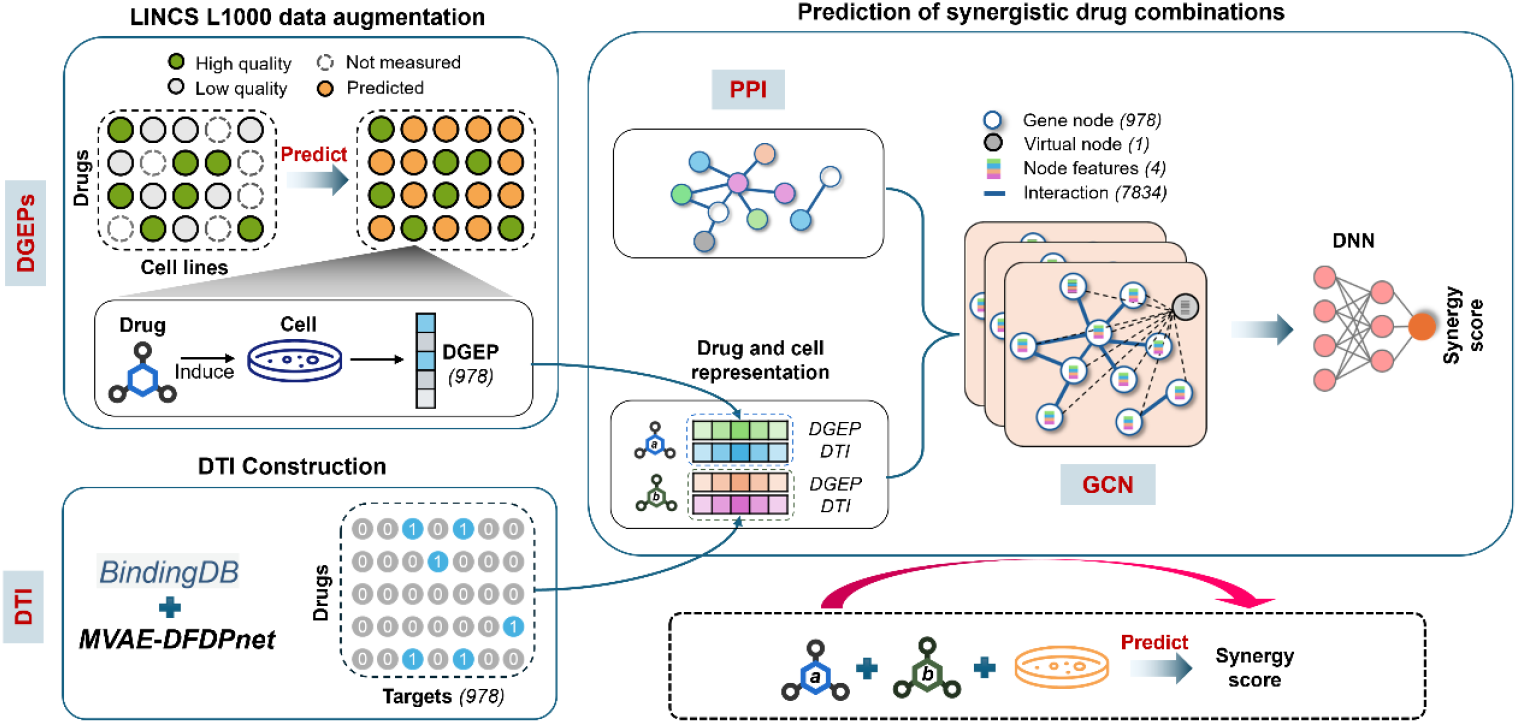
Overview of DRUGSYNC. In the first module, we employ high-quality DGEP data in LINCS L1000 project to impute and enhance lower quality measurements. In the second module, the DTI matrix was constructed utilizing data from MVE-DFDPnet DTI predictions, and the BindingDB database. In the third module, we construct the model specifically by building a GCN based on prior knowledge of PPI networks. Then, we utilize DGEP and DTI of two paired drugs as feature inputs to predict their synergistic scores within the DrugcombDB database.

### 2. Drug synergy prediction with augmented data

In the LINCS L1000 project, while the responses of compounds were tested across 161 cell lines, on average each compound was assayed in only about 5 cell lines. Notably, only 17.5% of the samples were classified as high quality, which required more than one biological replicate at Level 4 and a median Spearman correlation coefficient greater than 0.3 among the replicates. After training on the filtered high-quality data, our model achieved excellent performance. The overall median PCC between the predicted and experimental values of the model was 0.877 (Fig. 2A). The distribution of PCCs showed that the median PCCs for the top 10% and 50% were 0.967 and 0.925, respectively (Fig. 2B). These results demonstrate that our model can accurately predict drug-induced gene expression profiles, making it suitable for subsequent prediction tasks. It is noteworthy that the model’s performance may decline when applied to unseen drugs (PCC = 0.745) and cell lines (PCC = 0.694), as shown in Fig. 2C. After applying data augmentation techniques, the number of overlapping drug-drug-cell combinations between LINCS L1000 and DrugCombDB increased by 7-fold. This significant enhancement enriches the dataset, potentially improving the model’s ability to generalize and predict outcomes for novel compounds and cell lines.

**Fig. 2.**
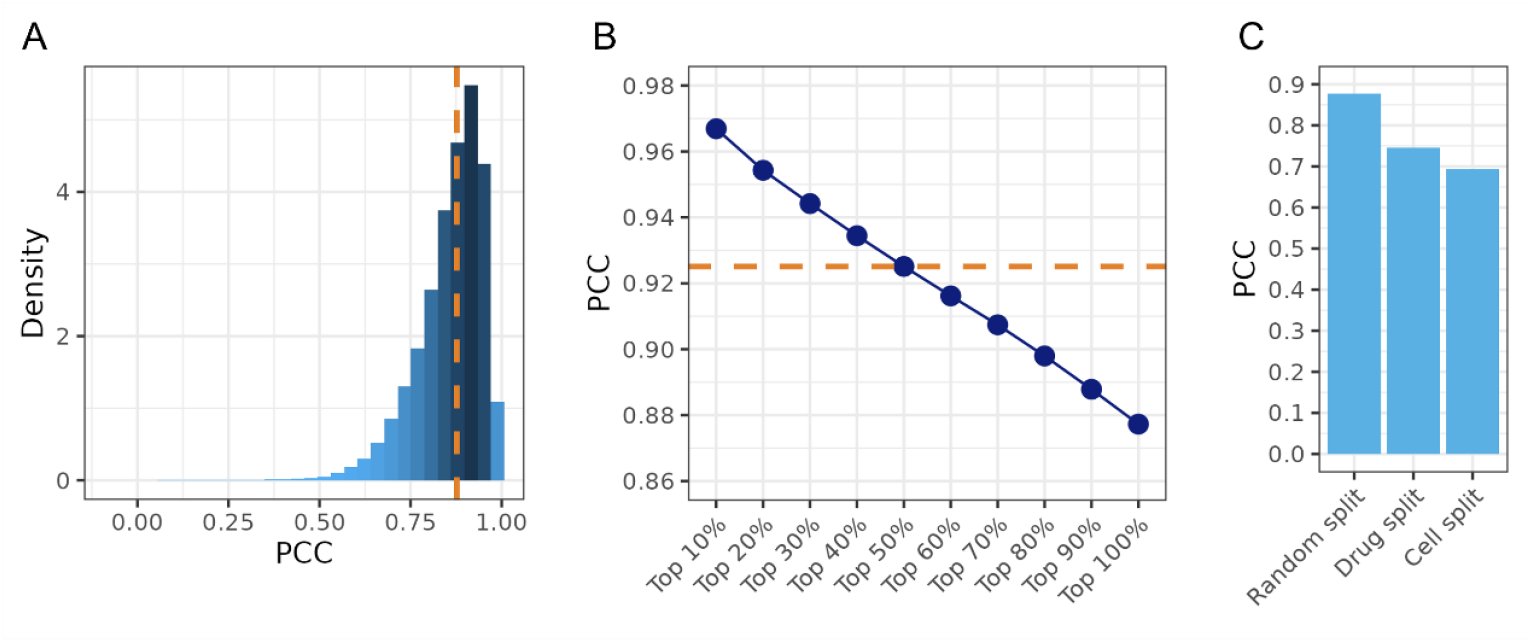
Performance of predicting the DGEP. (A) Distribution of the PCCs between the experimental and predicted sample. The orange line indicates the position of the median. (B) Cumulative plot for the average values of top x% PCCs. The orange line indicates the position of the top 50% PCCs. (C) The PCCs of MGEPs prediction model on new drugs (drug split) and new cell line (cell split) dataset.

### 3. DDS prediction with augmented data

We tested model performance on both regression and classification tasks with leave-combinations-out paradigm. When predicting the synergistic score, the model achieved a PCC of 0.76 and an R2 value of 0.33 (Fig. 3A). These metrics indicate a moderate positive correlation between the predicted and actual DDS scores. In synergy vs. non-synergy classification, the model demonstrated a BACC of 0.851 and a ROC-AUC of 0.934 (Fig. 3B).

**Fig. 3.**
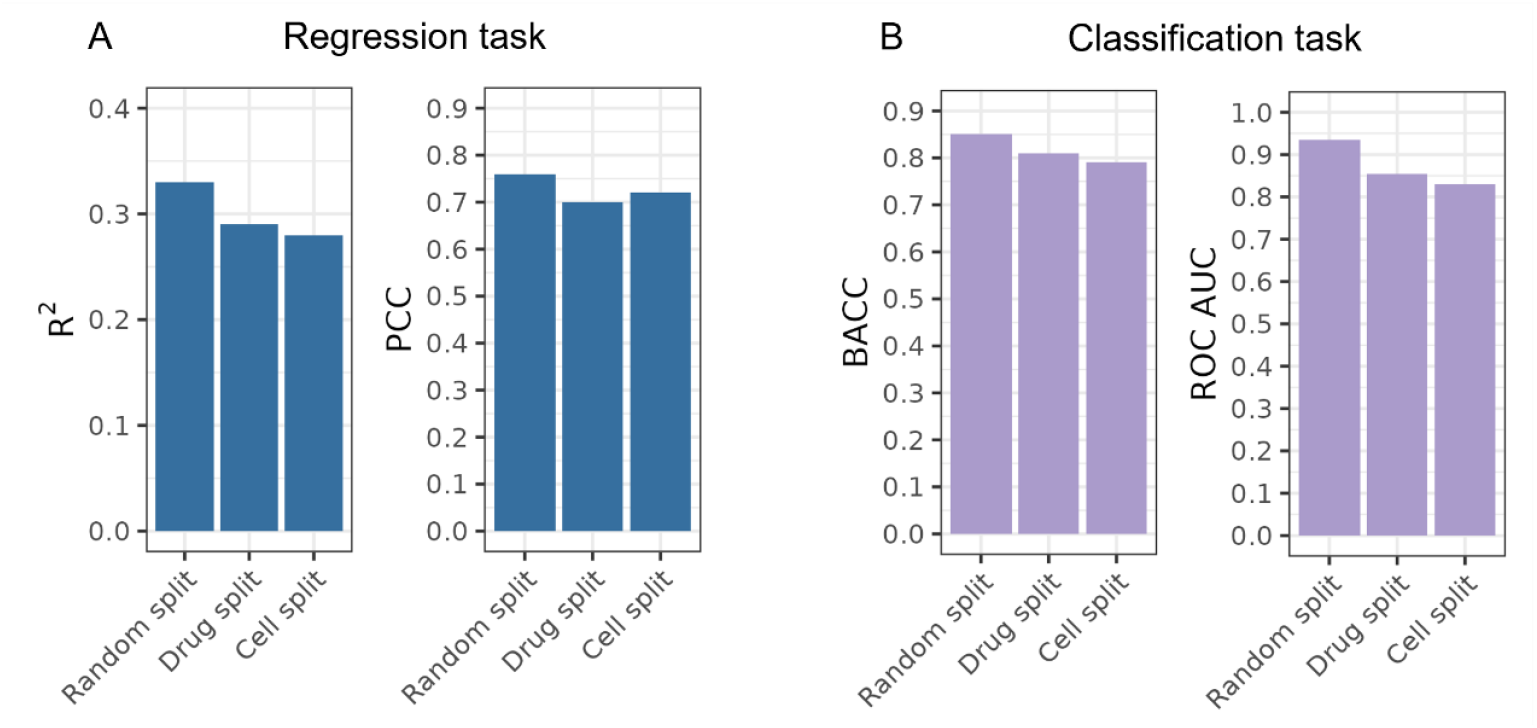
Performance of predicting the DDS. (A) Regression Task. We evaluated the model’s ability to predict the Bliss synergy score, using Pearson Correlation Coefficient (PCC, left) and R-squared (R^2^, right) as metrics. (B) Classification task. We tested the model’s performance on a classification task, where the labels were ‘synergistic’ and ‘non-synergistic’. The performance was assessed using Balanced Accuracy (BACC, left) and ROC Area Under the Curve (AUC, right).

Model generalizability was then tested with leave-drugs-out and leave-cells-out sample splitting. The results indicate that, even when evaluated on unseen drug or cell line, the model still maintains a high level of predictive accuracy. In regression tasks, the PCC under drug split and cell line split methods were 0.70 and 0.72, respectively (Fig. 3A). For classification tasks, the BACC achieved by the drug split and cell line split methods were 0.810 and 0.791, respectively. These findings underscore the generalizability of our method across different types of data splits (Fig. 3B).

### 4. Comparison with other prediction methods

To compare the performance of our proposed model against state-of-the-art models, we directly utilized the results reported in the DFFNDDS study [13], which were obtained by training on the DrugCombDB database. which were obtained on the DrugCombDB database. This approach allowed us to benchmark our model’s performance against established benchmarks using the same dataset, ensuring a valid and comparable evaluation. We selected models for comparison focusing on classification tasks, as these tasks are most relevant (Table 1). The results demonstrate that the performance of other models declined sharply using either leave-drugs-out or leave-cells-out method. In contrast, our model exhibits the least performance degradation using both methods, which highlights its superior generalizability on new data.

**Table 1.** Comparison with other prediction methods.

|  | Leave combinations out |  | Leave drugs out |  | Leave cells out |  |
| --- | --- | --- | --- | --- | --- | --- |
|  | BACC | ROC AUC | BACC | ROC AUC | BACC | ROC AUC |
| DRUGSYNC | **0.851** | **0.934** | **0.810** | **0.854** | **0.791** | **0.830** |
| #DFFNDDS [13] | 0.834 | 0.921 | 0.599 | 0.651 | 0.731 | 0.829 |
| #DeepDDS [12] | 0.784 | 0.900 | 0.586 | 0.621 | 0.713 | 0.812 |
| #DeepSynergy [11] | 0.762 | 0.860 | 0.588 | 0.630 | 0.629 | 0.630 |
| #MRGNN [30] | 0.737 | 0.845 | 0.557 | 0.574 | 0.692 | 0.806 |
| #GCNBMP [31] | 0.717 | 0.828 | 0.556 | 0.585 | 0.620 | 0.688 |
| #EPGCNDS [32] | 0.630 | 0.740 | 0.580 | 0.631 | 0.613 | 0.727 |
| #Match-Maker [16] | 0.638 | 0.746 | 0.586 | 0.649 | 0.650 | 0.701 |

### 5. Ablation study

To evaluate the contribution of different components to the overall performance of our framework, we conducted a series of ablation studies. These studies focused on the inclusion or exclusion of DTI features, the PPI network, and data augmentation (Fig 4A and B). In the presence of the PPI network, the model utilized GCNs to capture the complex relationships between genes. Without the PPI network, the model was reconfigured to a DNN. The following sections detail the results of removing or altering each component and their impact on the model’s accuracy. Specifically, we compared the DRUGSYNC results of: (i) DRUGSYNC without the PPI, (ii) DRUGSYNC without the DTI, (iii) DRUGSYNC without the pre-learned DGEP (i.e. with only experimental DGEP and much smaller sample size), (iv) DRUGSYNC without the PPI and DTI, (v) DRUGSYNC without PPI and pre-learned DGEP, (vi) DRUGSYNC without DTI and pre-learned DGEP, (vii) DRUGSYNC without PPI, DTI and pre-learned DGEP. We observed a significant decrease in BACC (from 0.851 to 0.652) when pre-learned DGEP were removed. Similarly, the removal of PPI network information led to a reduction in BACC (from 0.851 to 0.723). However, the removal of DPI resulted in only a slight decrease in accuracy. Overall, in the absence of a sufficient amount of data, the other components were not able to significantly enhance the model’s accuracy.

**Fig. 4.**
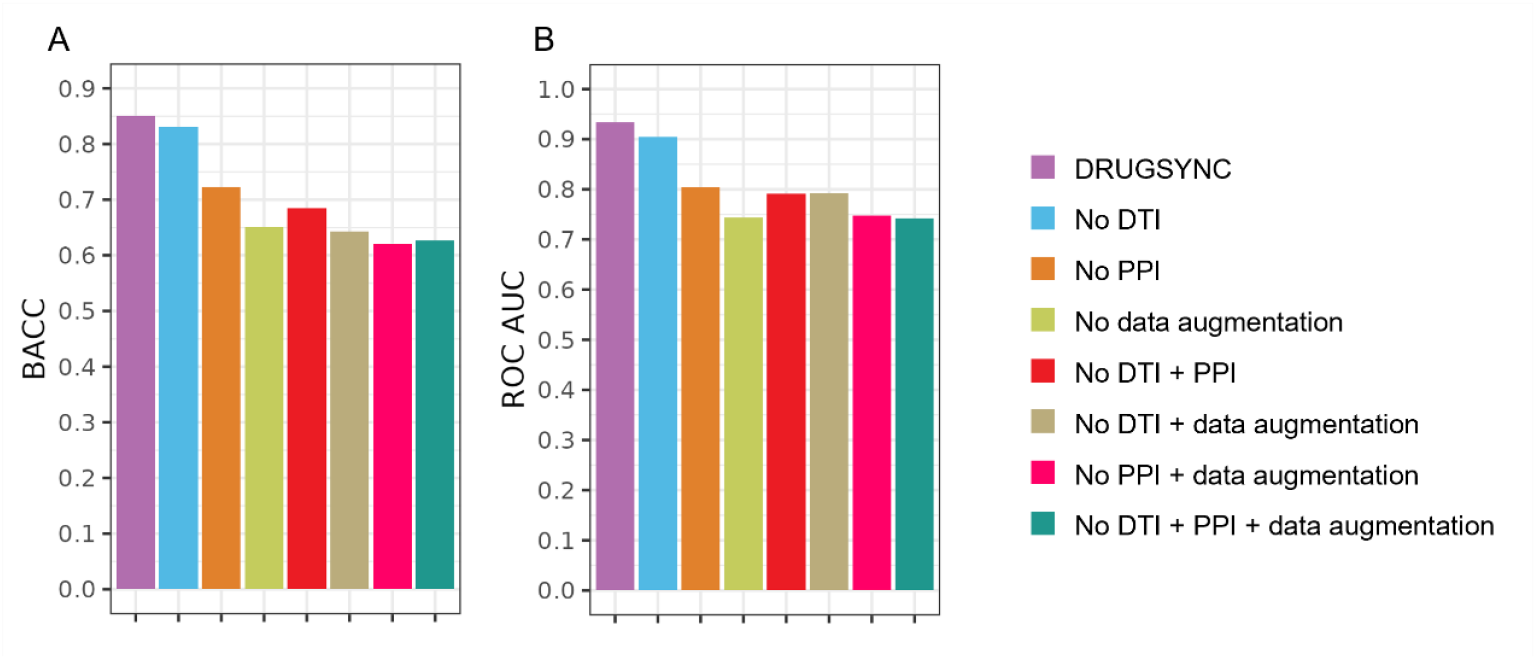
Impact of different model components on overall performance. This figure illustrates the effect of including or excluding various components (DTI features, PPI network, and data augmentation) on the overall performance of the model. The x-axis represents the different configurations, while the y-axis shows the BACC (A) and ROC-AUC (B).

## Discussion and conclusion

In this study, we introduced a novel framework, DRUGSYNC, for predicting synergistic drug combinations by integrating GCNs with pre-learned DGEP and DTI data. Our approach addresses the limitations of existing methods, particularly in terms of their performance on novel drugs and cell lines, by leveraging high-quality DGEP and incorporating prior biological knowledge through PPI networks.

Our approach offers several advantages over existing methodologies. By utilizing pre-learned DGEP and DTIs, we were able to capture the complex interplay between drugs and their cellular targets, which is often overlooked in models that focus solely on chemical descriptors. The incorporation of PPI networks into the GCN architecture allowed us to model the intricate network of genetic interactions, enhancing the model’s understanding of the underlying biological processes. This integration of multiple data types and the use of machine learning techniques contributed to the improved predictive power of our framework.

Previous studies have highlighted the importance of experimental DGEP, showing that the total contribution of DGEP features of 82.40% among others such as cell line GEP and molecular fingerprints according to an analysis of feature contributions in drug discovery tasks [20]. However, high quality experimental DGEP is sparse. While DGEP prediction is a way to expand training data, varying accuracy of predicted DGEP may impact its potential contribution to model performance. Our ablation studies also demonstrate that the use of high-quality, predicted DGEP substantially improves model performance.

Despite the promising results, there are areas for improvement. For example, our DGEP prediction module relies on the compounds present within the LINCS L1000. The limited size of the compound space significantly restricts our ability to investigate drugs not included in the LINCS L1000. In future work, we aim to utilize high-quality LINCS L1000 data to predict the DGEP with new compound solely based on structure information, thereby overcoming these limitations.

In conclusion, the DRUGSYNC framework represents a significant advancement in the field of DDS prediction. By effectively combining GCNs with high-quality DGEP and DTI data, our model not only achieves state-of-the-art performance but also demonstrates strong generalization capabilities.

## Methods

### Drug-induced gene expression profiles

We extracted cell line-specific DGEP data from the LINCS L1000 project [28]. We performed quality control on the data at Level 4, filtering out any datasets that had only a single biological replicate or those with multiple replicates but a median PCC less than 0.3 among them. For the data corresponding to various doses and time points of the same drug at level 5, we selected the 24-hour time point, which is the one with the most observations across all drug treatments. We then performed a weighted average of the different doses based on Pearson’s correlation coefficient, akin to the moderated z-score (MODZ) approach [28]. This consideration stems from previous reports indicating that the duration of drug exposure has a more significant impact on cellular gene expression profiles compared to dosage [23].

### Cell line gene expression profiles

We also apply gene expression profiles of untreated cell lines (without any drugs) to distinguish different cancer cell lines. The corresponding 978-dimensional gene expression profiles under vehicle control are extracted from the LINCS L1000 project as the feature data of cell lines. In LINCS L1000 project, vehicle control refers to the solvent used to administer compound treatments, which is usually DMSO.

### PPI

We downloaded the human PPI network from the STRING database v.12.0. (https://string-db.org/) [26,33]. To ensure the reliability of the interactions, only those with a STRING score > 700 were included in our analysis [26]. Due to the presence of isolated nodes and disconnected subnetworks within the dataset, an additional virtual node was introduced to facilitate global connectivity. The virtual node serves as a global scratch space, allowing information to travel long distances during the propagation phase [34]. Consequently, the final PPI network comprised 978 nodes and 7,834 edges.

### DTI

We have compiled drug-target information from the BindingDB database, that contains 2.9M data for 1.3M Compounds and 9.4K Targets [29]. Furthermore, we conducted additional predictions using our previously published MVAE-DFDPnet model, which has demonstrated superior DTI prediction capabilities [27]. The drug-target information was encoded into the GCN model as a binary node feature, where the value indicates the presence or absence of drug-target gene information.

### Drug synergy score

Drug Bliss synergy scores were downloaded from the DrugComDb (http://drugcombdb.denglab.org/main). In total, DrugCombDB includes 498,865 combination experiments, covering 5350 drugs and 104 cancer cell lines. We performed unique matching of drugs and cell lines across LINCS L1000 and DrugComDb using the SMILES notation for compounds and the COSMIC ID for cell lines. To account for variability across multiple replicate experiments, we adopted the median Bliss synergy score for each drug-drug-cell combination. Outliers were identified and excluded using the Interquartile Range (IQR) method, ensuring a robust statistical analysis of synergistic effects. For classification task, the Bliss synergy scores greater than 0 or less than 0 are considered “synergistic” or “non-synergistic” drug combinations, respectively.

### Build training and validation datasets

For paired data, validation on randomly split datasets often yields overly optimistic results. Besides the random split drug-drug-cell combinations (leave-combinations-out) into training and validation sets, it is also crucial to ensure that there is no overlap of any part of the paired data between the training and validation sets. Specifically, we employ drug split (leave-drugs-out) and cell split (leave-cells-out) methods to validate the model’s performance on novel drugs and cell lines. In the case of a leave-drugs-out, drugs present in the training set do not appear in the validation set. Similarly, with a leave-cells-out, cell lines used for training are distinct from those in the validation, ensuring that the model’s ability to generalize to unseen data is accurately assessed. This approach helps in obtaining a more realistic evaluation of the model’s predictive power on completely new data points. To evaluate the model’s performance and ensure its generalization capability, the dataset is split into 5 non-overlapping subsets. During each iteration, one of these subsets is used as the validation set, while the remaining four are combined to form the training set. This process is repeated five times, with each subset serving once as the validation set.

### Framework of the proposed model

The DGEP prediction model takes inputs in the form of triplet features (drug *i*, cell line *a* and cell line *b*). We define baseline gene expression profiles of cell line *a* and *b* as *G*_*a*_ and *G*_*b*_, respectively. Following exposure to drug *i*, the gene expression profiles of cell lines *a* are denoted by *G*_*ia*_. Each of these vectors *G*_*a*_, *G*_*b*_ and *G*_*ib*_ ∈ ℝ^*k*^, where *k* signifies the number of genes measured. The final feature vector for a given combination of drug *i*, cell line *a* and cell line *b* is constructed by concatenating these three components: *F*_*iab*_ = [*G*_*ia*_, *G*_*a*_, *G*_*b*_], where *F*_*iab*_ ∈ ℝ^3×*k*^ represents the concatenated feature vector. The constructed feature vectors *F*_*iab*_ used as inputs to a deep neural network (DNN) for training purposes. The objective was to predict the DGEP for cell line *b*, denoted as *G*_*ib*_ following treatment with drug *i*. The Root Mean Square Error (RMSE) is utilized as the loss function to quantify the discrepancy between the input and output.

For DDS prediction model. Given a PPI network, each node *v*_*m*_ in the graph represents a gene, and edges *e*_*mn*_ denote interactions between genes *v*_*m*_ and *v*_*n*_. For each gene *v*_*m*_, we associate a four-dimensional feature vector *X*_*m*_ = [*g*_*mi*_, *g*_*mj*_, *t*_*mi*,_ *t*_*mj*_ ]^*T*^, where: *g*_*mi*_, *g*_*mj*_ represent the expression levels of gene *v*_*m*_ under the influence of drug *i* and drug *j*, respectively. *t*_*mi*,_ *t*_*mj*_ are binary indicators (0 *or* 1) specifying whether drug *i* and drug *j* target gene *v*_*m*_. Additionally, each edge *e*_*mn*_ is attributed with a normalized STRING score *S*_*mn*_, which quantifies the confidence level of the interaction between genes *v*_*m*_ and *v*_*n*_. Thus, for a pair of drugs *i* and *j*, we construct a weighted graph *G* = (*V, E, S*), where: *V* is the set of nodes (genes) with associated feature vectors *X* = [*X*_1_, *X*_2_, …, *X*_*n*_]^*T*^, *E* is the set of edges indicating the interaction between genes, *S* is the set of normalized STRING scores for each edge, represented as a weight matrix *W*. The goal is to predict whether the drug *i* and drug *j* exhibits a synergistic effect. We formulate this as a binary classification problem, where the output is *y* ∈ {0, 1}, with *y* = 1 indicating synergy and *y* = 0 indicating no synergy. We employ a GCN to learn from both the graph structure and feature vectors. The GCN processes the graph *G* by aggregating information across neighboring nodes while considering the edge weights *W* that reflect the interaction confidence. Following three rounds of convolution, the output passes through three fully connected layers to produce a vector of 1 dimension. For classification tasks, we employ the Binary Cross-Entropy Loss as the loss function. For regression tasks, we utilize the RMSE as the loss function.

### Evaluation metrics

For the prediction of DGEP, we selected PCC, and R-squared (R^2^) as the metrics for the regression task. For the prediction of synergistic effects, which is a classification task, we used balanced accuracy score (BACC) and receiver operator characteristics curve (ROC) as the evaluation metrics.

### Generation and Aggregation of DGEP

If a drug has high-quality DGEP data for one cell line, it is possible to use our model to generate its DGEP for other cell lines. In cases where multiple sets of high-quality data are available, the approach would be to generate the target cell line’s data based on each of these datasets and then average the results.

